# Single-molecule insights into DNA gyrase in live bacteria

**DOI:** 10.64898/2026.06.21.733238

**Authors:** Aisha H. Syeda, Victoria A. Leek, Anthony Maxwell, Mark C. Leake

## Abstract

Molecular motors travelling along DNA introduce positive supercoils that present as barriers to replication leading to genome instability. To counter these, bacterial cells express DNA gyrase, a topoisomerase that introduces negative supercoils. While much is known about DNA gyrase from genetic and *in vitro* biochemical studies, the spatiotemporal dynamics of this enzyme remain a mystery. Only recently have we been able to observe the *in vivo* spatiotemporal dynamics down to single molecule level using advanced super-resolution microscopy techniques. We used Slimfield microscopy, a cutting-edge molecule microscopy technique to address the gap in our knowledge. We analysed a dual fluorescently labelled *Escherichia coli* strain expressing the replisome marker DnaN-mCherry along with mYPet-GyrB as the enzyme marker. We performed sequential Slimfield microscopy of the labelled proteins from the same strain and analysed *in vivo* GyrB dynamics in live *E. coli* cells in relation to the replisome. We find that the majority of replisomes are associated with GyrB. Inhibition of gyrase activity reduces the proportion of replisomes associated with GyrB. Interestingly, GyrB behaviour is distinct from that observed for GyrA in a previous study. Our results reveal the previously unknown dynamics of GyrB inside living bacterial cells highlighting the advantages of *in vivo* single molecule investigations. Our findings also demonstrate the importance of analysing all subunits of a functional enzyme complex to gain comprehensive understanding of its *in vivo* mechanisms. This study demonstrates the utility of single-molecule super-resolved microscopy as a valuable underpinning technology to understand *in vivo* behaviour of biomedically important molecules. Our insights will help impact discovery and development of novel antibiotics that interfere with gyrase function, thus contributing to tackling the growing problem of antimicrobial resistance.

## Introduction

Movement of molecular motors such as the replisome and RNA polymerase generates positive supercoils on DNA that present barriers to progression of these molecular motors. Hence timely removal of these supercoils is crucial for accurate motor function and genome stability. Inability to do so can lead to genome instability ultimately leading to cell death. Thus, interference with mechanisms that resolve positive supercoils is a valuable strategy to kill pathogenic microbes. In bacteria, DNA gyrase, a type IIA topoisomerase actively introduces negative supercoils to counter the accumulation of positive supercoils during translocation of molecular motors (McKie et al., 2021). This makes DNA gyrase a medically attractive antimicrobial target as it is found in bacteria (Nöllmann et al., 2007) but not in human cells. Indeed, the fluoroquinolone and aminocoumarin classes of antibiotics interfere with gyrase action leading to cell death (Bush et al., 2020; Gellert et al., 1976). However, widespread mutations in gyrase to counter these antibiotics adds to the growing problem of antibiotic resistance (Bush et al., 2020; Gellert et al., 1977; Hooper & Jacoby, 2016).

Our current level of understanding about gyrase activity in its native cellular environment is not sufficient to counter the mechanisms of its antibiotic resistance. The impact of different classes of antibiotics on the enzyme subunits *in vivo* is not fully understood as most of our understanding comes from traditional *in vivo* genetic, and *in vitro* approaches such as biochemical analyses and largely static structural biology “snapshots”. These techniques have their unique advantages in identifying factors involved in a biological function by *in vivo* mutant analysis and then their characterisation *in vitro* respectively. However, they are limited in that they provide a simplistic ensemble overview of protein/enzyme function while blurring out the behaviour of individual molecules in their natural cellular environment. The discovery and adaptation of fluorescent proteins to track molecules *in vivo* now enables us to probe the function of individual proteins in their native cellular environment (Payne-Dwyer et al., 2022; Reyes-Lamothe et al., 2010; Syeda et al., 2019; A. J. M. Wollman et al., 2024). Having a better understanding of the effect of antibiotics on gyrase activity and thereby on the processes they impact will allow us to identify mechanisms of antibiotic resistance. This will help us tackle the growing problem of antimicrobial resistance by enabling better informed combination therapies and novel antimicrobial development.

Slimfield microscopy (Plank et al., 2009) is a powerful technique that allows tracking of individual molecules to reveal key features of molecular processes, which has been used to gain transformative insights into a range of biological processes in live single bacterial cells involving DNA replication and remodelling (Badrinarayanan et al., 2012; Reyes-Lamothe et al., 2010; Syeda et al., 2019; A. J. M. Wollman et al., 2024), biomolecular condensate formation (Jin et al., 2021; Pei et al., 2025), gene expression in single budding yeast (A. J. Wollman et al., 2017), as well as following immune response signalling in multicellular tissue sections (Cosgrove et al., 2020; Miller et al., 2018), epigenetics signalling in whole root tips (Payne-Dwyer et al., 2025), and modified for in vitro single molecule imaging of DNA topology (Payne-Dwyer et al., 2025). It differs from standard epifluorescence in that the laser is focused on a narrow field of view of the sample (A. J. M. Wollman & Leake, 2016), producing a confocal excitation volume with up to 1000-fold high laser intensity. This results in high signal-to-noise ratio (A. J. Wollman & Leake, 2015) enabling millisecond and sub-millisecond timeframe sampling of individual molecules thus allowing us to infer the spatio-temporal behaviour of proteins in their native environment. Importantly, it allows super-resolved localisation of fluorescent proteins to a few tens of nanometre precision (Miller et al., 2017; Shepherd et al., 2021; A. J. Wollman & Leake, 2015) enabling us to identify individual enzyme behaviours beyond what is inferred using traditional approaches.

We used Slimfield to visualise the *in vivo* behaviour of the GyrB subunit of DNA gyrase. A previous study that analysed GyrA behaviour *in vivo* in relation to the replisome revealed that about 80% of GyrA clusters colocalised to the replisome (Stracy et al., 2019). To gain further mechanistic insights about the roles of the two gyrase subunits at the replisome in this study we used a dual labelled strain carrying mYPet-GyrB as the gyrase component along with DnaN-mCherry encoding the beta clamp of the replisome. Our analysis revealed that the *in vivo* copy number of GyrB is higher than that previously reported for GyrA. We find that 53% of GyrB is associated with 70% of DnaN. This association is less than that observed for GyrA in the previous study. Further, we were able to detect the disruption of this association upon treatment with gyrase targeting antibiotics ciprofloxacin or coumermycin A1 resulting in reduced localisation of GyrB with DnaN.

Combining results from a previous study, our results uncover different *in vivo* behaviours of the two gyrase subunits. Our results highlight the importance of comprehensive analysis of various subunits of a multi-subunit enzyme complex to gain novel insights regarding its *in vivo* behaviour.

## Materials and methods

### Purification of mYPet-GyrB

BL21(DE3) cells possessing pET28-MHL-mYPet-GyrB were grown to OD_600_ = 0.8 and induced with 0.8 mM IPTG. Cells were lysed using a Cell Disruptor and then centrifuged at 18000 rpm (1 hour). Purification of the supernatant was achieved using an Äkta Pure chromatography system, based upon a method previously described (Maxwell & Howells, 1999). Briefly, mYPet-GyrB was bound to a HisTrap column and eluted using with a gradient of 5-500 mM imidazole buffer containing 50 mM Tris pH 7.5, 10% glycerol, 2mM β-mercaptoethanol and 450 mM NaCl. Fractions containing mYPet-GyrB were dialysed overnight in TGED buffer (50 mM Tris-HCl pH 7.5, 10% glycerol, 1 mM EDTA, 1 mM DTT). TGED buffer was used to bind the protein to a MonoQ column, before elution with a gradient of TGED to TGED + 1 M NaCl.

### *In vitro* supercoiling assays

Supercoiling assays were conducted to test the *in vitro* activity of the fluorescent constructs compared to wild type. Time course assays were performed to study the effect of varied incubation periods at 37°C on the supercoiling activity. Relaxed DNA (pBR322), at a final concentration of 0.02 mg/mL, was used as a substrate for the supercoiling reactions. Assays were carried out in 35 mM Tris·HCl (pH 7.5), 24 mM KCl, 4 mM MgCl_2_, 2 mM DTT, 1.8 mM spermidine, 1 mM ATP, 6.5% (w/v) glycerol, 0.1 mg/mL albumin and volumes of dilution buffer and H_2_O were adjusted according to the volumes of gyrase subunits added to achieve a total volume of 30 μL. The supercoiling reactions were stopped by addition of 15 μL 2x STEB (40% sucrose, 100 mM Tris pH 8, 100 mM EDTA, 0.5 mg/mL bromophenol blue), followed by 24:1 chloroform isoamyl alcohol (30 μL) to isolate DNA into the aqueous phase. Samples were vortexed, centrifuged at 13200 rpm (2 minutes) and then ∼15 μL of the aqueous phase was analysed on a 1% agarose gel. Gels were run at 80-100V in TAE (40 mM Tris base, 20 mM glacial acetic acid and 1 mM Na_2_EDTA), stained with 1 μg/mL ethidium bromide (30 minutes), destained in TAE buffer and finally visualised under UV light.

### Slimfield microscopy

A Slimfield fluorescence microscope was used that had been developed around a Zeiss microscope body using a Nikon CFI SR HP Apo TIRF 100XC Oil objective lens and an *xyz* nano positioning stage (Nanodrive, Mad City Labs). The microscope utilised narrow epifluorescence excitation of 7 μm full width at half maximum (FWHM) in the sample plane to generate Slimfield illumination from fluorescence excitation of 514 nm and 561 nm wavelength lasers passed through a ∼3 Keplerian beam de-expander. Imaging was performed via a custom-made colour splitter utilizing a dual-pass green/red dichroic mirror centered at long-pass wavelength 560 nm and emission filters with 25 nm bandwidths centered at 542 and 594 nm (Chroma Technology Corp., Rockingham, Vermont, USA) onto a Photometrics 95B CMOS camera, magnified to 50 nm/pixel. Initially, two frames of brightfield were acquired to image the cell boundary. The fluorescence images were then sequentially acquired in the red and yellow channels. The samples were first excited with 20 mW of 561 nm laser until bleached (200 frames) and images were recorded for the mCherry channel. Thereafter, mYPet images were acquired by exciting with 10 mW of 514 nm laser for 1000 frames. Both brightfield and fluorescence acquisitions were performed at maximum gain at 5 ms/frame.

### Preparing samples and obtaining fluorescence data

The *E. coli mYPet-gyrB dnaN-mCherry* dual labelled strain was streaked onto an LB plate and incubated at 37°C overnight. Single colonies from the plate were then inoculated into LB media and grown at 37°C shaking at 180 rpm for 5 hrs. The culture was diluted (1:50) into M9 supplemented with 0.2% glycerol as carbon source and grown overnight at 37°C to A_600_ of 0.4–0.6. Next morning, the cultures were diluted 10 times into fresh M9 glycerol and grown again to A_600_ of 0.4–0.6. Cells were centrifuged and immobilised for imaging on a glass slide (Reyes-Lamothe et al., 2010). Briefly, gene frames (Life Technologies) were attached on a glass microscope slide to form a well. Then 500 µl of M9 media with 1% agarose was added to the well to form a thin layer. After the pad was dry, small droplets of the *E. coli* culture were pipetted onto the pad and allowed to dry. The well was then covered with a plasma-cleaned glass coverslip for imaging. Where required, cells were incubated with 10 μg/ml ciprofloxacin or 100 μg/ml coumermycin A1 for 30 min prior to slide preparation.

### Analysing the data

Slimfield images can be analysed using in-house generated custom MATLAB code and Python codes (Miller et al., 2015; Shepherd et al., 2021; A. J. Wollman et al., 2015). The signal-to-noise ratio was set to >0.4. The intensities of single mYPet and mCherry molecules were determined from stepwise photobleaching analysis of foci that were overtracked beyond their bleaching point. These single molecule intensities were then used to calculate the stoichiometry of fluorescent foci by simply dividing the intensity of the foci with the determined intensity of single fluorophores. Colocalisation between fluorescent foci was established if the positions of tracks from two different channels were within a 256 nm window (Stracy et al., 2019). Significance testing for colocalisation was performed using Fisher’s exact tests. Odds ratios were calculated as the ratio of the odds of the tracks in a channel from the treated sample to the that in the untreated sample.

## Results and discussion

### Results

#### mYPet-GyrB is catalytically active

Purified mYPet-GyrB was tested for supercoiling activity and compared against purified native untagged GyrB. The final concentrations of GyrA and GyrB were 2.1 nM and 50 nM respectively. We note the requirement of higher concentrations of GyrB in our *in vitro* assays as compared to GyrA. The reactions containing supercoiled plasmid DNA as template were incubated on a time course ranging from 0 to 60 minutes. The reactions were then run on 1% agarose gels to qualitatively assess the supercoiling activity of GyrB (figure 1). The supercoiling activity of mYPet-GyrB was comparable to native wild-type GyrB in our assays, indicating that the fusion is fully functional. We note that after longer incubation periods a linear band became more prevalent with the mYPet-GyrB.

**Figure 1:**
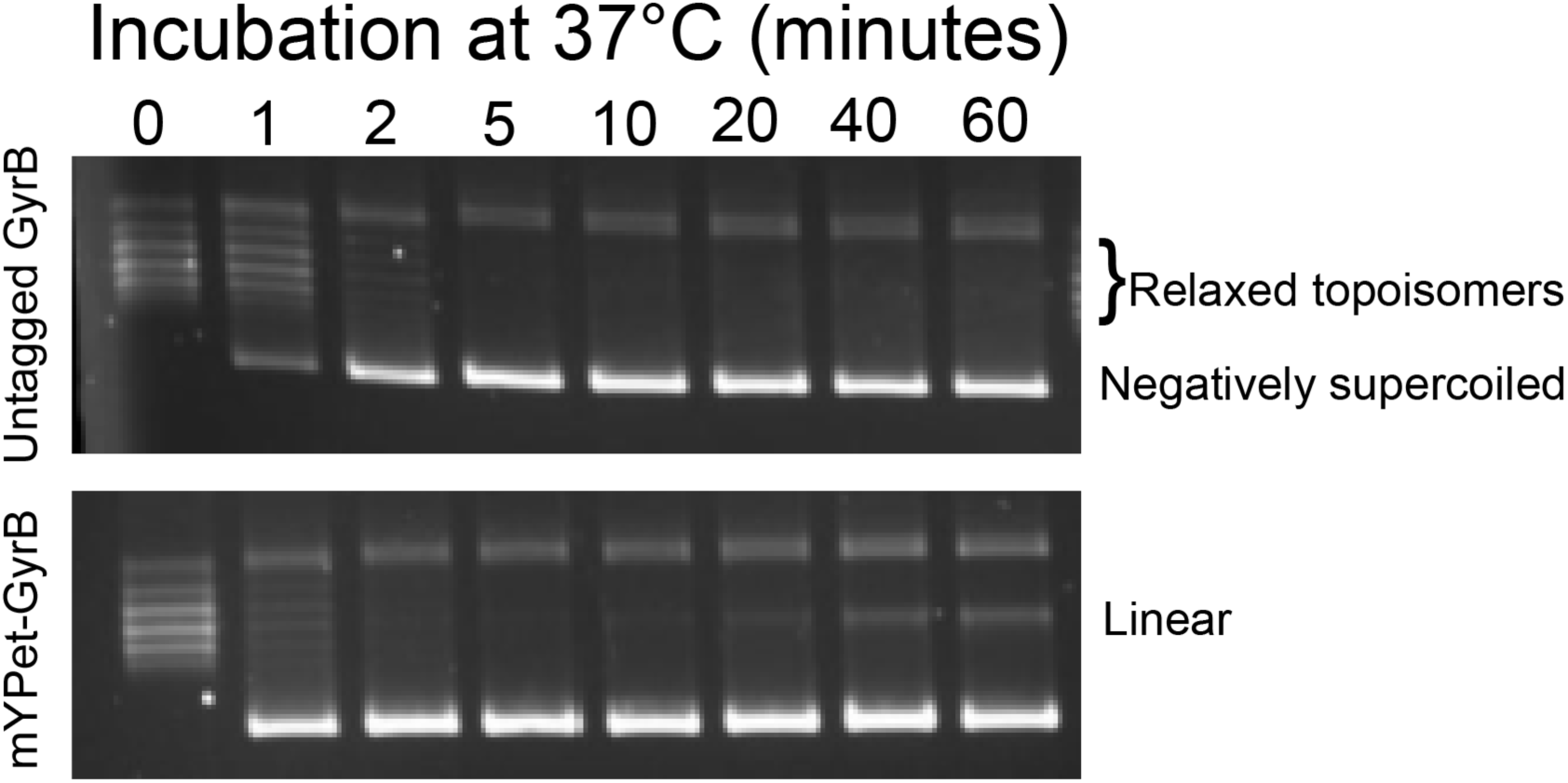
mYPet-GyrB is catalytically active. Gel-based supercoiling assays were performed with GyrA in combination with native untagged GyrB (top panel) and mYPet-GyrB (bottom panel). Time-course supercoiling assays were performed with fixed concentrations of both subunits and checked on a 1% agarose gel. Lane 0 contains relaxed topoisomers at the beginning of the assay before initiation of enzyme activity. The subsequent lanes are labelled with the corresponding number of minutes of incubation with the enzyme. The gels show that by about 5 minutes both labelled and unlabelled GyrB containing reactions catalyse complete conversion of the open circular topoisomers to negatively supercoiled forms.

#### Slimfield enables millisecond sampling and single molecule detection *in vivo* allowing quantification and localisation behaviour of GyrB

To study the *in vivo* dynamics of *E. coli* GyrB inside living cells, we utilised a dual-labelled fluorescent strain comprising mYPet-GyrB and the replisome marker DnaN-mCherry (Reyes-Lamothe et al., 2010). Using Slimfield microscopy, we observed the relationship between the replisome and GyrB in real-time using high speed sequential laser excitation. The dual labelled DnaN-mCherry mYPet-GyrB strain was sequentially imaged and images from individual channels were recorded for each cell. A schematic of the Slimfield setup is illustrated in figure 2. The captured images were then analysed using bespoke analysis code (Miller et al., 2015) to reveal their stoichiometry and colocalisation dynamics in real-time.

**Figure 2:**
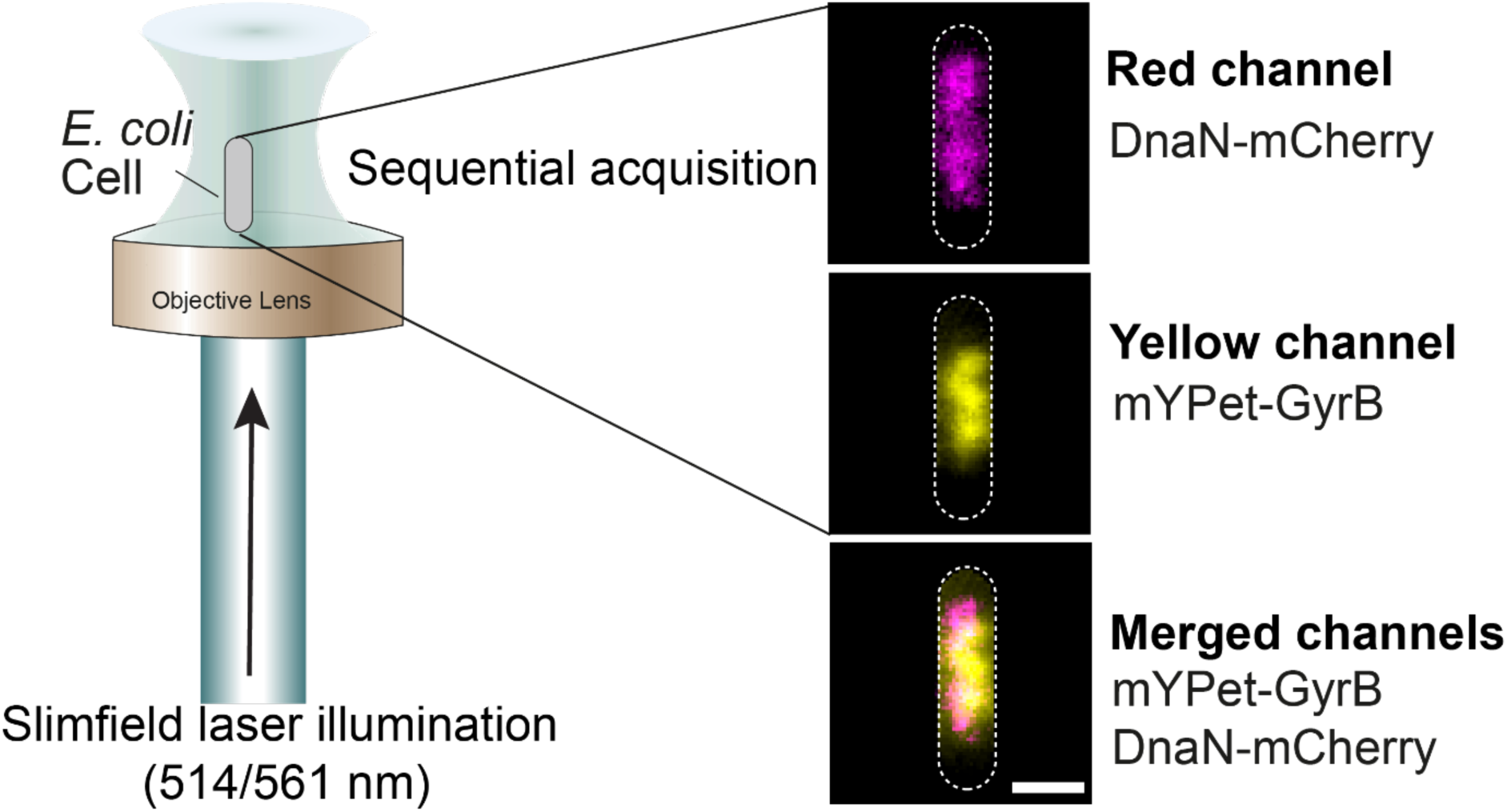
Simplified Slimfield schematic. Dual colour Slimfield setup that enables single-molecule tracking in live bacterial cells. DnaN-mCherry was imaged by illuminating the samples with a 561 nm laser (magenta) and mYPet-GyrB was imaged using a 514 nm wavelength laser (yellow). Cell outline is indicated by dashed outline. The images were acquired sequentially. The relative *in vivo* positions of fluorophores from both channels are illustrated in the bottom panel showing merged channels. Horizontal bar: 1 μm.

We quantified the copy number of mYPet-GyrB from the fluorescent data to assess expression of GyrB *in vivo*. We observed that GyrB is expressed at an average of 1670 ±39 molecules ranging from 756-4630 molecules per cell (figure 3a).

**Figure 3:**
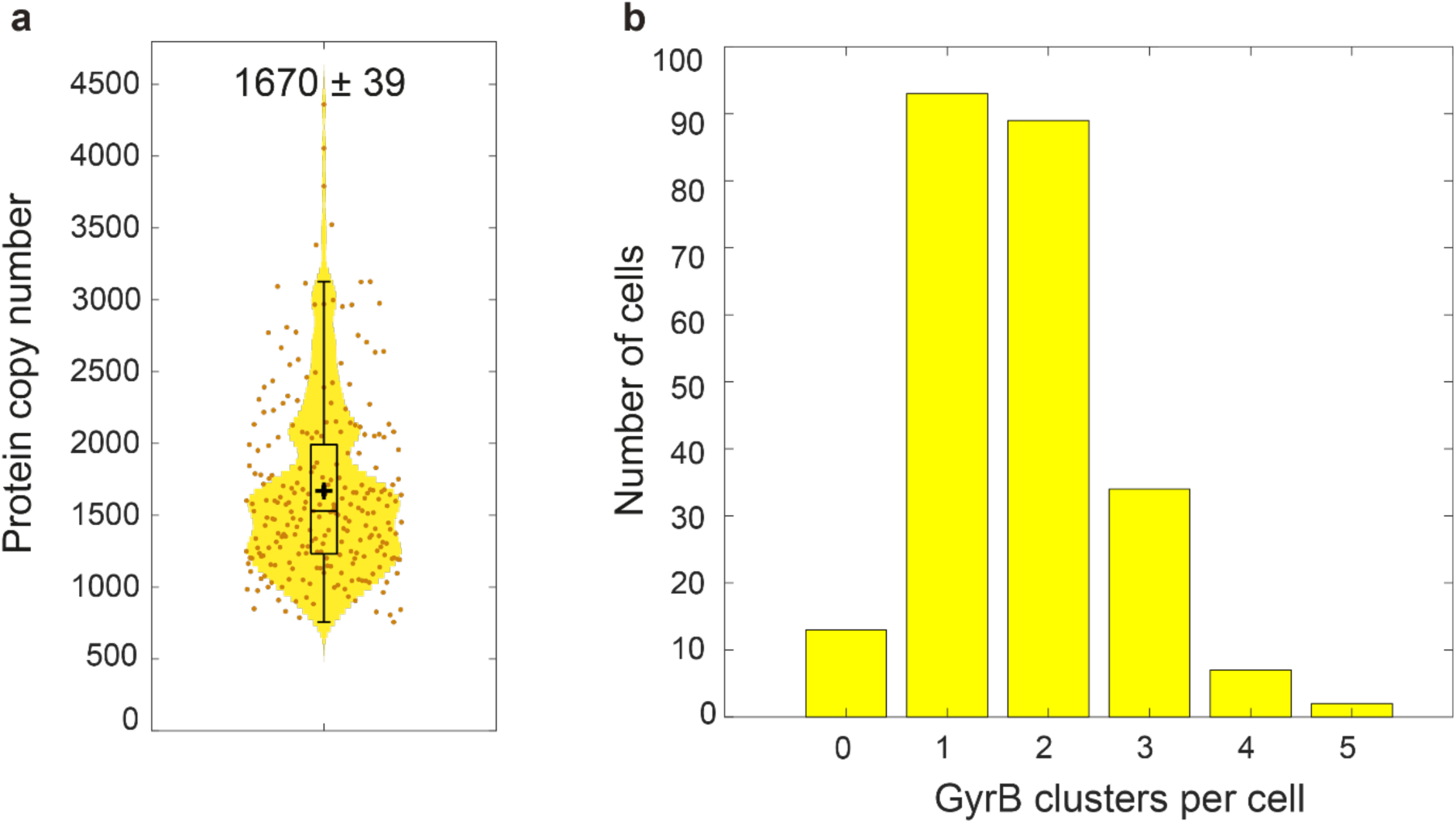
**a.** Violin plot distribution of GyrB copy numbers estimated from Slimfield data. Each data point indicates the estimated copy number from a single cell. The average copy number is indicated by the + symbol and the horizontal bar indicates the median. **b.** Histogram depiction of distribution of GyrB clusters per cell. Most cells had 1-3 clusters.

We observed clustering of GyrB into distinct clusters in 94.5% cells corresponding to an average of 1.73 ± 0.06 distinct clusters per cell. From these data we also infer most cells had one or two clusters with a substantial proportion also having three clusters (figure 3b). The average number of molecules in clusters per cell were determined to be 64.7 ± 1.63. From the average number of clusters and molecules in clusters per cell, a simple multiplication of the values therefore inferred the number of molecules in clusters per cell to be 112 ± 3 molecules. From the calculated average copy number of 1670 ± 39 per cell, this represents a very small proportion of molecules in clusters of only 6.7 ± 0.2 % molecules. This value indicates that the sampling time of 5 ms may be slow causing the molecules to blur and artificially underestimate the number of molecules in clusters. Future experiments with a higher sampling rate such as 2 ms may provide a more precise estimate of molecules in clusters.

#### GyrB forms multi-subunit clusters with a dimeric periodicity of GyrB

We observed distinct clusters corresponding to GyrB throughout the cell. We quantified GyrB stoichiometries using a counting approach that utilises stepwise photobleaching of the observed fluorescent proteins (Stracy et al., 2019). Kernel density estimations suggest a predominantly dimeric spacing of GyrB as observed by association of two molecules within 40 nm (figure 4a, inset showing zoom-in). Additionally, spacing in multiples of two was also observed. This observation agrees with biochemical estimations of two molecules of GyrB in a gyrase heterotetramer (Klevan & Wang, 1980; Sugino et al., 1980). Surprisingly, GyrB distribution with an average of 38 ± 1 molecules per cluster indicated that the clusters contain about 19-20 GyrB dimers.

**Figure 4:**
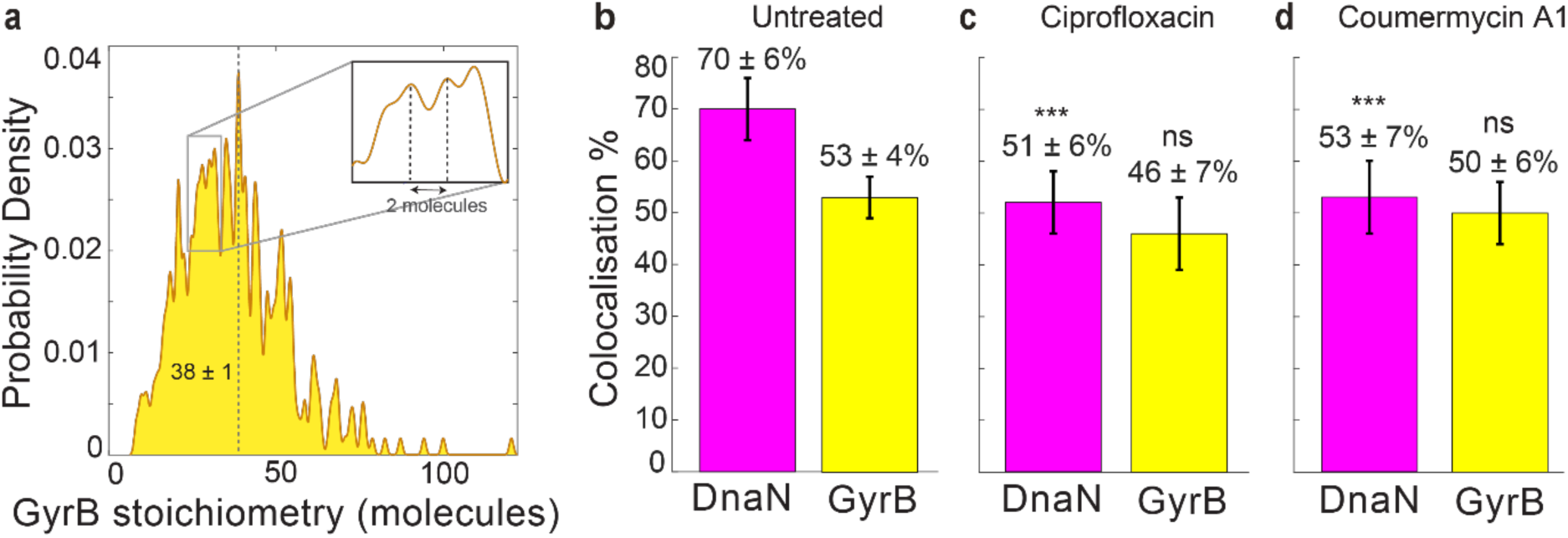
Dual colour sequential Slimfield reveals clustering of GyrB and its colocalisation behaviour with DnaN. **a.** GyrB stoichiometries shown as kernel density estimations (Llorente-Garcia et al., 2014). Intervals between peaks indicate the number of molecules present in a single molecular complex *in vivo*. The dashed vertical line indicates average stoichiometry. Inset shows a zoomed-in view to visualise a spacing interval of two molecules. **b - d.** Bar charts with error bars (s.e.m) depicting DnaN clusters colocalised to GyrB (magenta) and GyrB clusters colocalised to DnaN (yellow) in untreated **(b)**, ciprofloxacin-treated **(c)** and coumermycin A1-treated **(d)** cells. Fisher’s exact tests were performed for significance by comparing against the untreated; *** = highly significant (P<10^-3^), ns = not significant (P>0.05).

#### GyrB colocalises with the majority of replisomes and this colocalisation is disrupted by gyrase targeting antibiotics

We also analysed the tracked data (Miller et al., 2015) to determine the colocalisation of DnaN and GyrB. We found that 53 ± 4% of GyrB tracks were colocalised to 70 ± 6% DnaN tracks (figure 4b). We then treated the cells with gyrase inhibiting antibiotics to observe if this changes the colocalisation behaviour of GyrB to the replisome. The fluoroquinolone ciprofloxacin interferes with GyrA catalysis by lodging itself in the cleavage intermediate, thus preventing religation of the cleaved DNA strands (Bush et al., 2020; Hooper & Jacoby, 2016). The aminocoumarin coumermycin A1 interferes with GyrB ATP hydrolysis by competitively binding into the ATP binding pocket of GyrB (Sugino et al., 1978). We found that treatment with ciprofloxacin significantly reduced the colocalisation of DnaN and GyrB. 46 ± 7% of GyrB tracks (odds ratio = 0.77, p = 0.17) were colocalised with only 46 ± 7% of DnaN tracks (odds of colocalisation = 0.46, p = 0.2 X 10^-3^) (figure 4c). Similarly, we found coumermycin A1 to also reduce colocalisation of GyrB with 50 ± 6% tracks (odds of colocalisation = 0.91, p = 0.58) to only 53 ± 7% of DnaN tracks (odds ratio = 0.5, p = 1 X 10^-3^) (figure 4d).

## Discussion

We used a dual labelled strain expressing mYPet-GyrB and DnaN-mCherry to understand the behaviour of GyrB with relation to the replisome in its native cellular environment and also to understand its dynamics in the context of known GyrA behaviour from a previous study (Stracy et al., 2019). We found that GyrB is expressed at 1670 ± 39 copies per cell. GyrA was reported earlier to express at 1450 ± 550 copies per cell (Stracy et al., 2019). This unexpected observation reveals differential expression of GyrA and GyrB *in vivo*. An interesting biological observation in our study was that GyrB forms clusters with 38 ± 1 molecules per cluster. This is in excess of that calculated for GyrA at 24 ± 2 molecules per cluster (Stracy et al., 2019). Our results therefore reveal a higher and non-stoichiometric clustering of GyrB molecules than that reported for GyrA. Interestingly we also noted a higher requirement of GyrB in our *in vitro* assays as compared to GyrA, lending support to our finding.

We also observed GyrB colocalised to 70 ±6% DnaN. This is lower than that observed for GyrA in an earlier study where about 80% of GyrA tracks were colocalised to DnaN (Stracy et al., 2019). Whether these differences in expression and clustering are a result of experimental variation or a bona fide observation will be obvious in future investigations by directly comparing the two subunits. The colocalisation of GyrB with DnaN was significantly reduced upon treatment with ciprofloxacin and coumermycin A1.

Analysing a single subunit of a multisubunit complex only provides partial insight into the *in vivo* behaviour of an enzyme. Unlike biochemical investigations that require all components of a multisubunit complex, single molecule approaches allow analysis of behaviour of a single subunit. While this is an advantage as it allows one to probe the natural *in vivo* environment of an enzyme, an inherent danger is the extrapolation of its behaviour to all other subunits within the enzyme. Our study serves as a paradigm for analysing the various subunits that make up a macromolecular assembly to get a comprehensive understanding of enzyme activity by unmasking the behaviour of individual components.

A caveat to our current approach is that it suffers from artefacts due to comparison of observed GyrB activity in one strain to GyrA activity in another strain. These artefacts are compounded by differences in growth conditions and imaging platforms. We also note that in the current study we performed sequential acquisition of fluorescence signal that may result in inaccuracy in localisation determination. An ideal future study involving both subunits labelled in the same strain with alternating laser excitation would provide further mechanistic insights by eliminating artefacts due to strain, growth conditions and differences in microscopy setup.

## Acknowledgements

We thank Ji-Eun Lee for technical support in generating the Slimfield datasets used in this study. We also thank Pawel Zawadzki for the gift of *mYPet-gyrB dnaN-mCherry E. coli* strain. AS was funded from BBSRC grant BB/W000555/1. ML was funded from EPSRC grant EP/Y000501/1 and BBSRC grant BB/R001235/1. A.M. was supported by the BBSRC funded Institute Strategic Programme Harnessing Biosynthesis for Sustainable Food and Health (HBio) (BB/X01097X/1) and a Wellcome Trust Investigator Award (110072/Z/15/Z).

